# Evidence for Alternative Complement Cascade Activation in Primary CNS Vasculitis

**DOI:** 10.1101/329862

**Authors:** Caleigh Mandel-Brehm, Hanna Retallack, Giselle M. Knudsen, Alex Yamana, Rula A. Hajj-Ali, Leonard H. Calabrese, Tarik Tihan, Hannah A. Sample, Kelsey C. Zorn, Mark P. Gorman, Jennifer Madan Cohen, Antoine G. Sreih, Jacqueline F. Marcus, S. Andrew Josephson, Vanja C. Douglas, Jeffrey M. Gelfand, Michael R. Wilson, Joseph L. DeRisi

**Affiliations:** Department of Biochemistry and Biophysics, University of California, San Francisco, CA, USA; Department of Pharmaceutical Chemistry, University of California, San Francisco, CA, USA; Department of Rheumatology/Immunology, Cleveland Clinic, Cleveland, OH, USA; Department of Pathology and Laboratory Medicine, University of California, San Francisco, CA, USA; Department of Neurology, Boston Children’s Hospital, Boston, MA, USA; Division of Neurology, Connecticut Children’s Medical Center, Hartford, CT, USA; Division of Rheumatology, University of Pennsylvania, Philadelphia, PA, USA; Kaiser Permanente, San Francisco Medical Center, San Francisco, CA, USA; UCSF Weill Institute for Neurosciences, San Francisco, CA, USA; Department of Neurology, University of California, San Francisco, CA, USA; Chan Zuckerberg Biohub, San Francisco, CA, USA

## Abstract

The central nervous system (CNS) has a dedicated network of blood vessels to support the physiological activity of the brain, spinal cord and meninges. Consequently, inflammation of CNS vasculature can have devastating effects on neurological function. A lack of understanding regarding the molecular pathology of CNS vasculitis impedes the development of better diagnostics and effective therapies. Here, we analyze the proteome of cerebrospinal fluid from patients with biopsy-confirmed Primary Angiitis of the Central Nervous System (PACNS) relative to non-inflammatory control patients and patients with Reversible Cerebral Vasoconstrictive Syndrome (RCVS), a syndrome that clinically mimics PACNS in several aspects. In PACNS, we find significant elevation of apolipoproteins, immunoglobulins and complement cascade components. Notably, we find a bias towards activation of the alternative complement pathway with elevated levels of the terminal cascade component, complement C5. Given the recent treatment successes of Anti-Neutrophil Cytoplasmic Antibody (ANCA) vasculitis with the C5 receptor inhibitor, CCX168 (Avacopan), our results suggest that complement C5 inhibitors may also prove useful as therapeutic interventions for PACNS.

## Introduction

The molecular basis of blood vessel inflammation, or vasculitis (also referred to as angiitis), is poorly understood. This gap in knowledge is exemplified by Primary Angiitis of the Central Nervous System (PACNS), an inflammatory disease affecting the blood vessel walls in the brain, spinal cord and meninges^1^. Without treatment, PACNS is frequently progressive and fatal. While PACNS usually presents in adulthood, it can also occur in children^2,3^. Treatment paradigms utilizing broad immunosuppressants including corticosteroids, cyclophosphamide, and mycophenolate mofetil, based largely on the results of therapeutic trials of various types of systemic vasculitis, have been adapted for use in PACNS. While these treatments have improved the outcome of PACNS, they have adverse side-effects, and nearly half of patients relapse^4^. A lack of basic research in this area remains an obstacle for improving PACNS therapeutics and served as the motivation for this study.

Molecular analysis of cerebrospinal fluid (CSF) is a compelling approach for understanding PACNS pathophysiology. CSF is an accessible biologic fluid that circulates throughout the cerebral ventricular system and bathes neural tissue contained within the meninges and parameningeal structures of the brain and spinal cord. Molecular analysis of CSF can provide diagnostic information regarding disease pathologies occurring within these structures^5^. Imaging abnormalities in PACNS are common and often show evidence of inflammation within the meninges and parenchyma of the CNS^2,6^. In addition, CSF protein concentration is elevated in ∼80-90% of PACNS cases^5,7^. To our knowledge, there has been no systematic study to interrogate PACNS CSF for specific protein abnormalities that may shed light on the chronic inflammatory pathophysiology of the disease and/or provide molecular targets for therapeutic and diagnostic utility.

Here, we report a proteomic analysis comparing the CSF profiles of biopsy-proven PACNS patients to the CSF profiles from Non-Inflammatory Controls (NIC) and controls with Reversible Cerebral Vasoconstriction Syndrome (RCVS) that, early in the disease course, can mimic PACNS clinically and radiologically^8^. Compared to these control groups, we show that PACNS CSF exhibits molecular abnormalities that implicate immunoglobulins, the complement cascade pathway and lipoproteins in PACNS pathophysiology. More specifically, we observe significant alteration of several regulatory components of the alternative pathway, and elevated levels of the terminal cascade including complement C5 (C5), complement C8 (C8) and complement C9 (C9). In previous studies C5 has been shown to be a therapeutically relevant target for systemic Anti-Neutrophil Cytoplasmic Antibody (ANCA) vasculitis, and recent clinical trials of C5 inhibitors further support this notion^9^. Our results suggest a rationale for investigating whether C5 inhibitors could serve as a novel therapy for PACNS.

## Results and Discussion

### Study Design and Patient Cohort

A PACNS diagnosis is suspected when neurological deficits persist in the absence of systemic inflammation or evidence of infection and all potential alternative etiologies have been excluded. To confirm a PACNS diagnosis, the current gold standard involves histopathological analysis of blood vessel walls in a biopsy of the meninges, the brain parenchyma or both^1^. The histopathologic hallmarks of PACNS include evidence of transmural, lymphocytic predominant (minimal neutrophils) or granulomatous inflammation restricted to small to medium blood vessels in the CNS^10^. However, due to the “patchy” appearance of lesions and a pathological bias towards smaller blood vessels, biopsies often miss lesions and are only diagnostic in approximately 50-80% of cases^1,11^. For the purposes of this study, as a means of identifying direct manifestations of PACNS CNS pathophysiology, we restricted our CSF analysis to patients for whom a systemic vasculitis was not identified (e.g., no clinical evidence for systemic involvement, including serum ANCA antibody negative) with biopsy-proven PACNS (Figure 1A-F).

**Figure 1:**
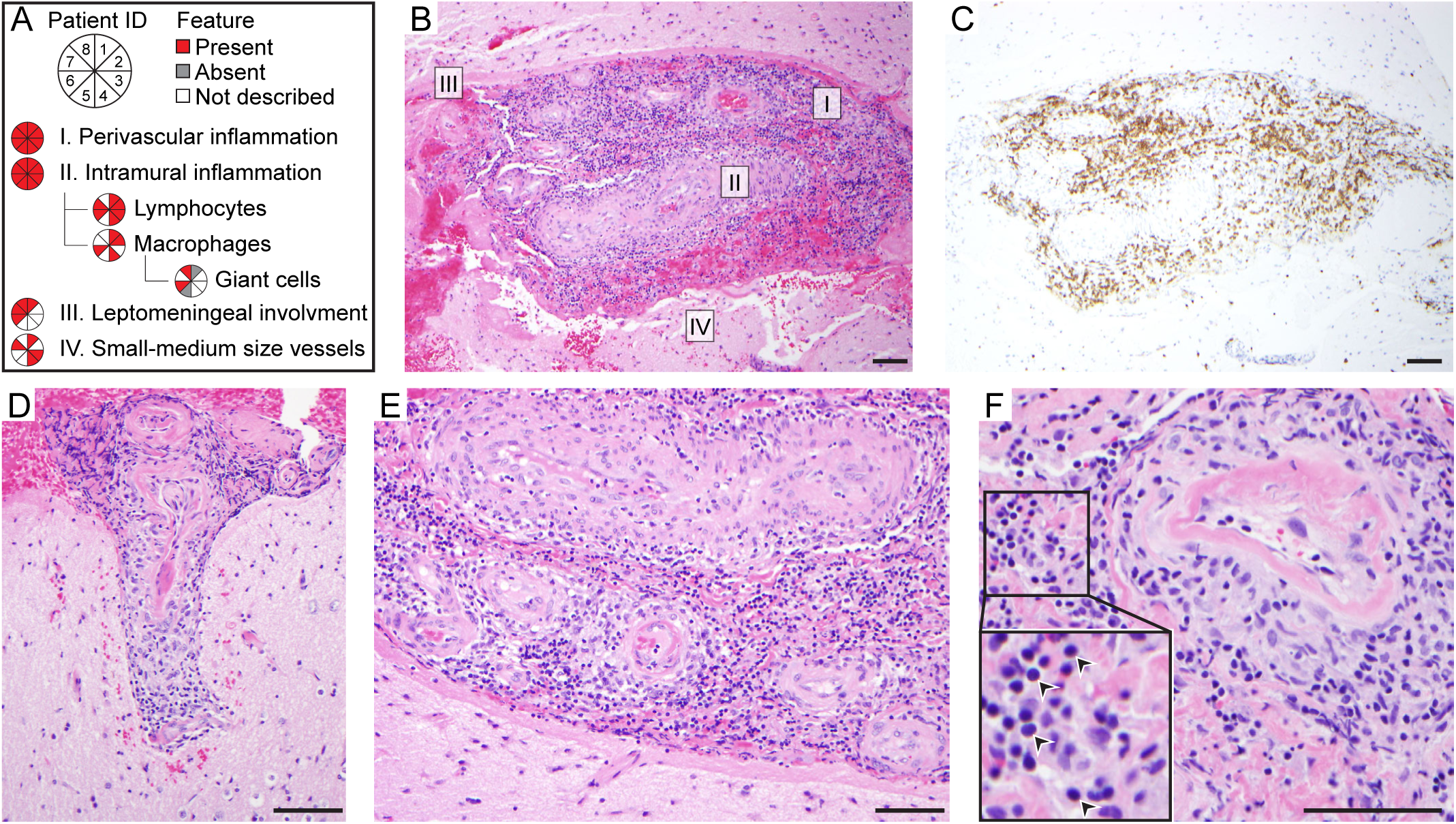
Histopathological features of biopsy proven PACNS cohort. (A) Summary of histopathological findings from the biopsy report of each PACNS patient used in this study (n = 8). All patient biopsies showed inflammation of CNS blood vessels. Immune cell infiltrates specifically enriched for T-cells were observed within (intramural) and around (perivascular) the blood vessel wall. (B-F) Representative images from a patient biopsy (ID:3) demonstrating several hallmark features of PACNS histopathology. Tissue sections were stained with H&E (B, D-F), or anti-CD3 antibodies (C) to identify T cells. Representative images for the histopathological findings described in (A) are indicated in panel (B). Arrowheads in the inset in panel (F) indicate lymphocytes. Scale bar, 100 µm.

The majority of our biopsy-proven PACNS patients were refractory to initial treatment strategies or required continual immunosuppression to achieve remission (Supplemental File S1). These observations reflect the chronicity of inflammation of the cerebral vasculature, which is a distinguishing feature of PACNS pathophysiology^10^. However, a straightforward analysis of PACNS CSF for molecular correlates of chronic inflammation is complicated by non-specific, acute cerebrovascular pathologies. For this reason, we analyzed an RCVS patient cohort, in addition to a set of NIC patients (Supplemental File S1). RCVS mimics some aspects of PACNS but exhibits only acute blood vessel constriction^8^. By directly comparing PACNS and RCVS patients, we sought to identify the molecule(s) or molecular pathways that contribute to the chronic inflammatory processes that define PACNS specific pathology.

### Mass Spectrometry analysis of CSF from NIC, RCVS and PACNS

For quantitative proteomic analysis, CSF protein from PACNS, NIC and RCVS cohorts was quantified by standard Bradford assay (Supplemental File S2) and five micrograms was subjected to quantitative mass spectrometric analysis (See Materials and Methods). Spectral datasets acquired from each sample were searched against the SwissProt Human Protein database to identify unique proteins. To report protein abundance, the total spectral count for each protein was normalized by the total number of spectral counts acquired per sample. A total of 1044 unique proteins were identified across NIC, PACNS and RCVS cohorts (Supplementary File S3). Using these data as input into an unbiased clustering analysis, we find that individuals in the NIC cohort are more similar to each other than to those in PACNS, indicating the NIC and PACNS cohorts form two distinct CSF proteomic profiles (Supplemental File S4).

Proteins in PACNS were considered differentially regulated from NIC controls if they had a statistically significant fold change of 1.5 or greater across two statistical tests of varying stringency (See Materials and Methods). Using this criterion, we identified 283 proteins that were differentially regulated in PACNS, with 61 up-regulated proteins and 222 down-regulated proteins compared to NIC (Supplemental File S5). Using the Database for Annotation, Visualization and Integrated Discovery (DAVID) Bioinformatics resource, we found that dysregulated proteins in PACNS showed significant enrichment for “Complement and Coagulation Cascades”, “Apolipoproteins”, “Immunoglobulins” and “Cell Adhesion Molecules” (Supplemental File S6).

To determine whether the altered proteins we identified in PACNS were due solely to blood brain barrier damage we utilized data from our RCVS patients. When compared to NIC controls, we found the correlates of acute blood vessel damage, including acute phase proteins and blood coagulation markers, among the most robustly elevated proteins in RCVS CSF. We found several Hemoglobin proteins (HBG1, HBB, HBA1, HBD), Serum Amyloid Apolipoproteins (SAA1, SAA2) and Carbonic Anhydrases (CA1, CA2) to be elevated in both the PACNS and RCVS cohorts (Figure 2). Notably, the common proteomic abnormalities in PACNS and RCVS were enriched among the coagulation component of the “Complement and Coagulation Cascade” identified by DAVID, whereas no dysregulated proteins were shared between PACNS and RCVS cohorts that belonged to the complement cascade. Thus, the significantly altered proteins within the complement pathway (properdin (CFP), Complement C4 Binding Protein A (C4BPA), Complement C4 Binding Protein B (C4BPB), Ficolin-3 (FCN3), Decay Accelerating Factor (CD55), MAC-inhibitory Protein (CD59), Carboxypeptidase N catalytic chain (CPN1), Carboxypeptidase N subunit 2 (CPN2), C5, C8, and C9) that occur in PACNS are independent of the non-inflammatory, cerebrovascular injury seen in RCVS. From these findings, we conclude that PACNS pathophysiology exhibits elements of damage to blood brain barrier but also exhibits specific changes in the complement cascade, apolipoproteins and immunoglobulins that cannot be explained by acute blood brain barrier damage alone.

**Figure 2:**
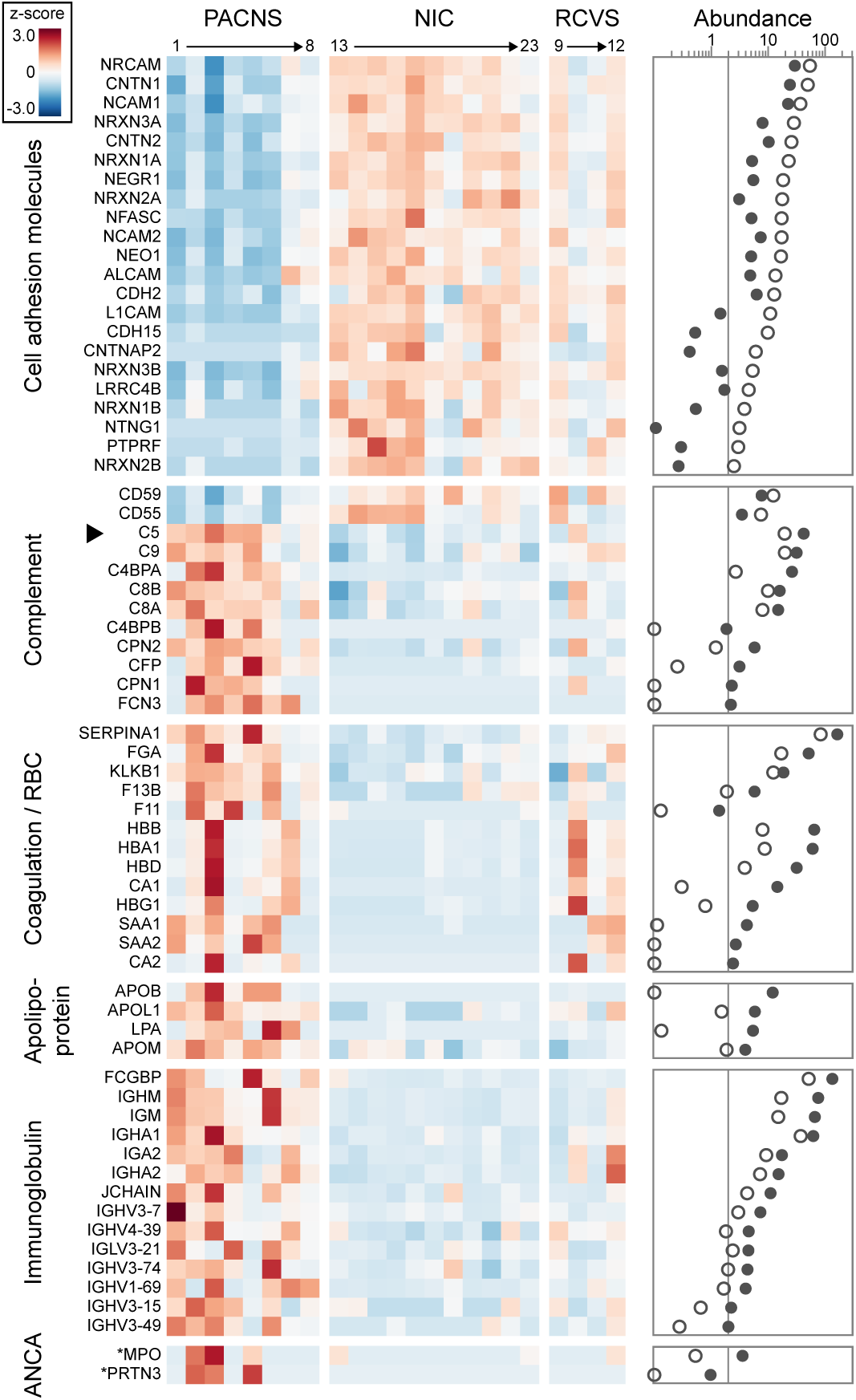
Regulatory proteins of the complement pathway, apolipoproteins, immunoglobulins and cell adhesions proteins among dysregulated proteins in PACNS CSF. The relative and absolute protein abundances for a subset of differentially regulated proteins in PACNS are reported. The subset of proteins was manually curated to reflect findings of significant clinical interest, with functional classifiers informed by DAVID and KEGG annotations. *(Heatmap, left)* Heatmap displays relative protein abundance across individual samples (PACNS, n=8, NIC, n=11, and RCVS, n=4). Values plotted in each mini-box represent the z-score of the normalized spectral counts. Note specificity and reproducibility of molecular findings pertaining to the complement cascade in PACNS. *(Scatter plot, right)* The mean protein abundance for each protein is reported for NIC (open circles) and PACNS (closed circles) cohorts. Values are plotted on a log scale, vertical line denotes an abundance of 2 normalized spectral counts to highlight minimally expressed proteins (abundance < 2 normalized spectral counts). Note the robust increase in abundance for a specific subset of proteins within the complement cascade that normally have minimal to no expression in NIC. Differential expression was evaluated using DESeq2 and t-test (see Materials and Methods), with significance defined as having fold change >1.5 and BH-adjusted p <0.05 for both tests. Asterisk denotes proteins that do not meet criteria for statistical significance but are of clinical interest.

As indicated above, apolipoproteins, specifically ApolipoproteinB100 (ApoB100) and lipoprotein(a) (LPA), and immunoglobulins (IgA, IgM) were robustly altered in the PACNS patients. These proteins are of particular interest as ApoB100 and LPA have been implicated in the chronic inflammatory process associated with atherosclerosis^12^. For example, evidence of a complex immune response to ApoB100, mediated by T-cells and IgM antibodies, is commonly observed in the progressive development of atherosclerotic lesions. A protective role of anti-ApoB100 IgM antibodies has been suggested, although not rigorously evaluated. Interestingly, we also observed an increase in IgM in PACNS and speculate that elevated IgM in PACNS CSF could be a compensatory mechanism in PACNS as well.

Transmembrane Cell Adhesion Molecules (CAMs) expressed in neural tissue were also enriched among the 221 downregulated proteins in PACNS (Supplemental S7). We localized the recovered peptides from mass spectrometry according to protein domain annotations assigned by Uniprot and found that PACNS CSF exhibited robust down-regulation in the extracellular domains of transmembrane proteins, normally present in NIC (Supplemental S8). These changes may occur at the transcriptional or post-transcriptional levels, or represent abnormal proteolytic processing of extracellular domains. An intriguing possibility is that loss of the ectodomains is linked directly or indirectly to the gain of complement and apolipoproteins. Regardless, we anticipate that these data will provide a useful resource as CSF analyses accumulate from patients with other types of neurological disorders.

Lastly, in the CSF of several (3 of 8) PACNS patients we identified Myeloperoxidase (MPO) and Proteinase 3 (PRTN3, or PR3), two previously described antigens in ANCA associated vasculitis^13^–^16^. These ANCA-associated antigens are expressed in the granules of neutrophils and monocytes and have ascribed roles in active inflammation. Similar to PACNS, ANCA-associated vasculitis exhibits chronic inflammation and destruction of blood vessel walls, however, the disease more frequently targets the kidney, gastrointestinal tract and lungs, rather than the CNS^17^. As described earlier, all our PACNS patients’ serum ANCA antibody test was negative and none of our patients’ vasculitis was identified by routine clinical testing to involve organs outside the CNS. Thus, we speculate that elevated levels of MPO and PRTN3 may be present in both types of vasculitis; however, the secondary antibody response to these proteins may manifest only in the context of systemic inflammation (i.e., ANCA-associated vasculitis).

### The “Complement Phenotype” of PACNS

Deregulation of the complement cascade pathway in PACNS is the most robust and clinically significant molecular finding of this study. These findings are highly reproducible across PACNS patients, constituting a “Complement Phenotype,” which predicts a hyperactive alternative complement pathway and elevated signaling through the terminal cascade (Figure 3).

**Figure 3:**
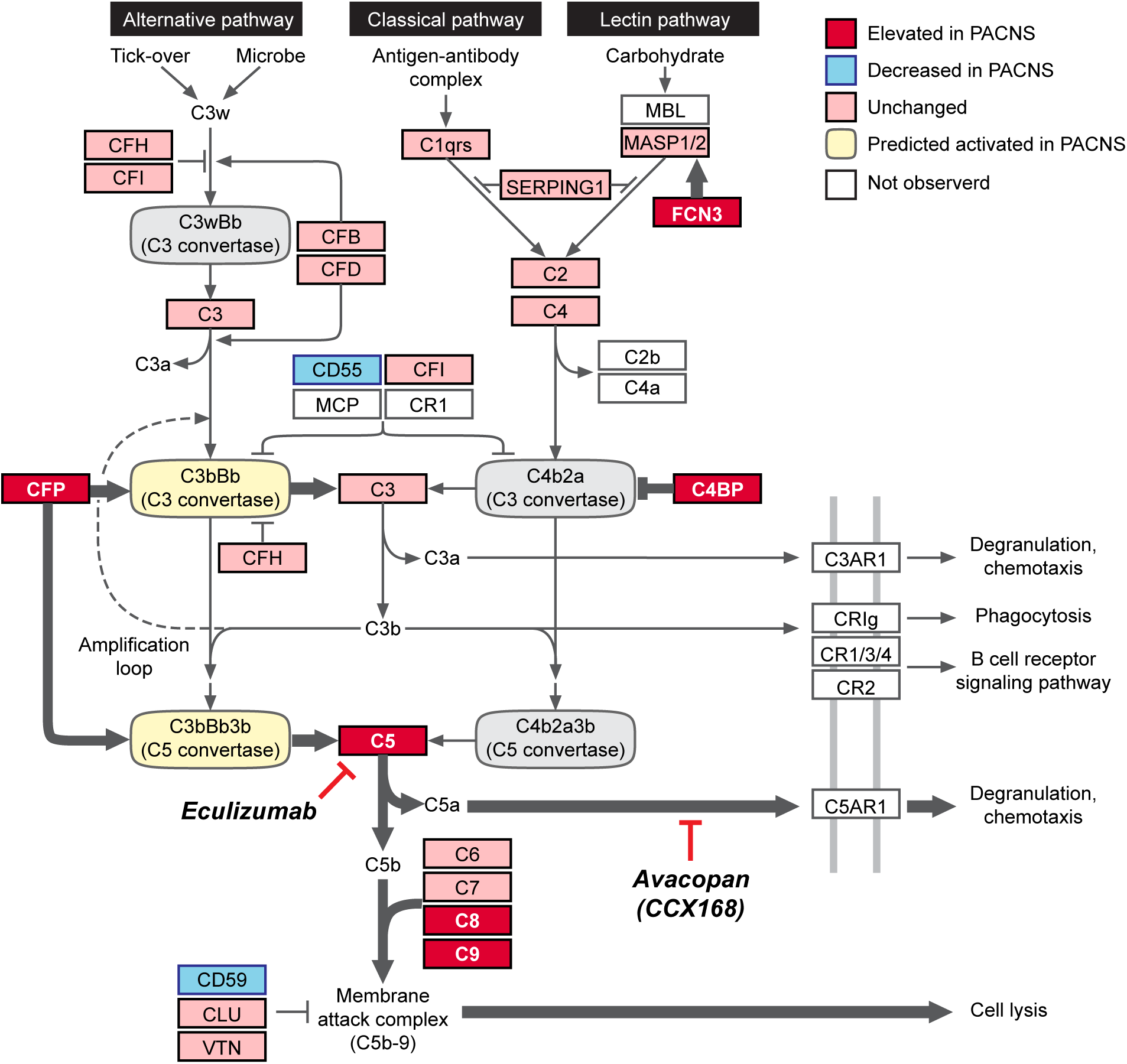
Proposed model of the “Complement Phenotype” of PACNS. The activity profile for the complement cascade in PACNS CSF is informed by proteomic comparison of CSF between PACNS and NIC cohorts (Figure 2, supplemental S5). Molecular data collected in Figure 2 is overlaid onto the complement cascade pathway (adapted from KEGG, hsa04610). The fold change in protein abundance between PACNS and NIC was evaluated for all proteins annotated in the pathway. Proteins are reported as Elevated (red, >1.5 fold significant increase in PACNS), Unchanged (pink, not significantly changed in PACNS), Decreased (blue, >1.5 fold significant reduction in PACNS) or Not Observed (white, no abundance in NIC or PACNS). The prediction for C3 and C5 convertase activity (Activated, yellow; Inhibited, gray) is informed specifically by molecular changes observed in the complement regulatory proteins CFP, C4BP, CD59 and CD55. The proposed model predicts a shift towards activation of the alternative pathway (CFP, CD55), active inhibition of Classical Pathway (C4BPA and C4BPB) and elevated signaling from the terminal cascade (CD59).

The complement cascade is comprised of two separate pathways, the Classical Pathway (homologous to Lectin Pathway) and the Alternative Pathway^18^. These pathways are molecularly unique at the level of the complement C3 (C3) and C5 convertases that drive cleavage of C3 and C5, respectively. In PACNS CSF, the bias towards activation of the alternative pathway is largely driven by elevated protein levels of CFP (properdin), the only known fluid phase positive regulator of the alternative pathway^19^. More specifically, CFP stabilizes the alternative pathway convertases, which are otherwise extremely unstable and easily deactivated. The bias towards the alternative pathway is also evident in the elevated protein levels of C4BPA and C4BPB, well-studied inactivators of the classical pathway specific C3 convertases^20^. Altogether, there appears to be concomitant *activation* of alternative pathway specific C3 and C5 convertases and *inhibition* of classical pathway C3 convertases. Thus, net activity favors the alternative pathway over the classical pathway.

Downstream of the alternative pathway, we observed evidence of an increase in C5, elevated protein levels of the MAC complex (C8A, C8B, C9) and a reduction in CD59, an inhibitor of the MAC complex^18^. Taken together, these protein changes are consistent with the activation and stabilization of the alternative pathway resulting in increased terminal cascade signaling. The increase in terminal cascade signaling may result from sustained alternative pathway signaling, including increased secretion of CFP from neutrophils, but could also result from dysfunction within suppressive factors such as negative feedback regulators.

Our finding of elevated levels of C5 is of particular significance given the recent success of a clinical trial using CCX168 (Avacopan) to treat ANCA-associated vasculitis^9,21^.CCX168 inhibits C5 signaling by binding to the C5a receptor (C5aR1 / CD88). Our findings, together with the recent data on CCX168, reveal a previously unappreciated overlap in pathophysiology between PACNS and ANCA-associated vasculitis.

### Orthogonal validation of elevated complement C5 in CSF of PACNS patients using commercially available ELISA

To assess levels of complement C5 independent of mass spectrometry, we employed a commercially available C5 ELISA assay (LS-bio, LS-F24462-1). Currently, C5 levels are routinely tested by ELISA in serum but not in CSF. Regardless, C5 was readily detectable in PACNS CSF by ELISA for 5 of 8 patients. In contrast, C5 signals were below the level of detection in all NIC and RCVS CSF control samples (Figure 4). Thus, ELISA-based measurements of C5 were consistent with the mass spectrometric analysis of CSF, confirming elevated C5 in PACNS CSF.

**Figure 4:**
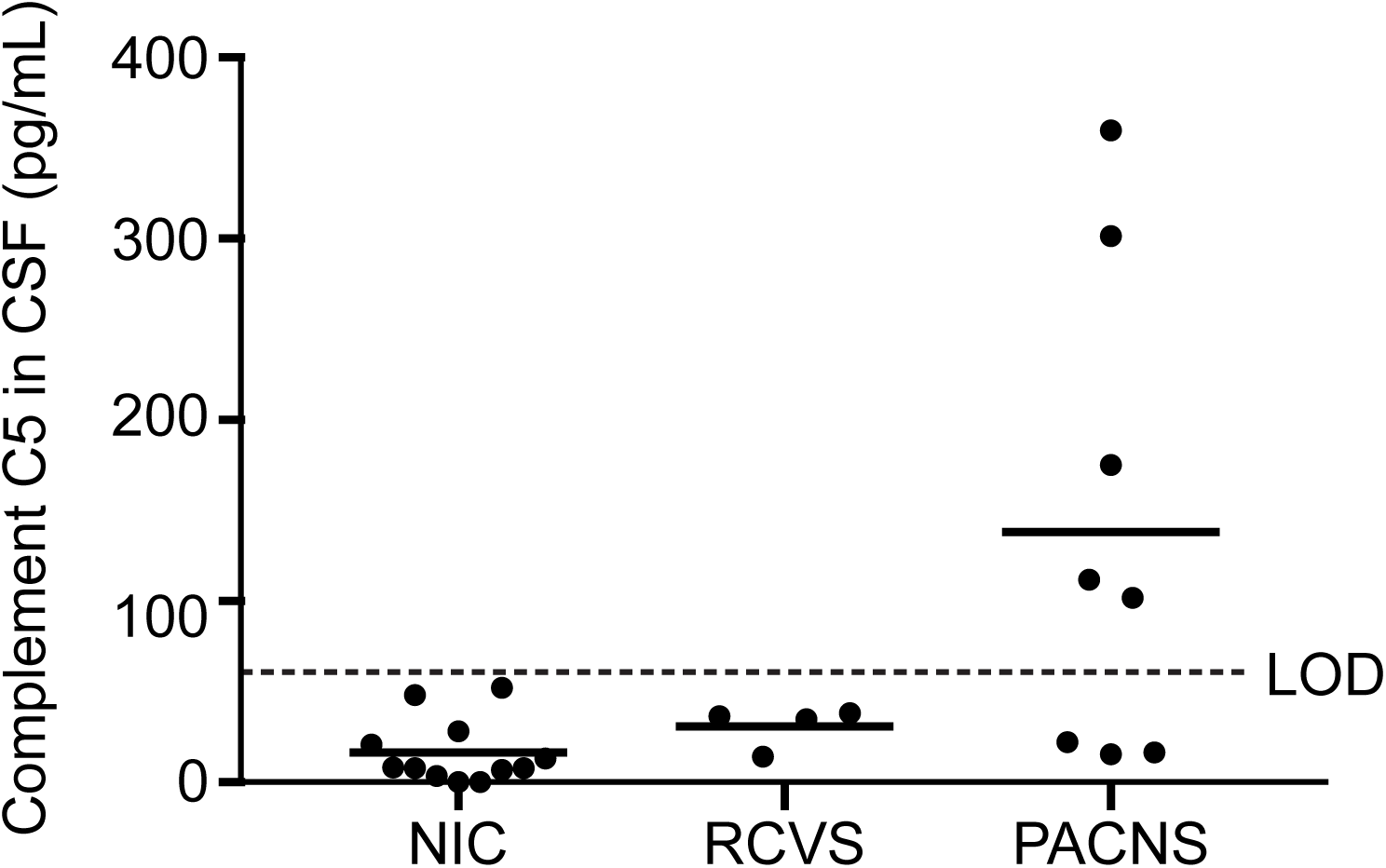
Orthogonal validation of elevated C5 in CSF of PACNS patients using commercially available ELISA. The concentration of C5 (pg/mL) in the CSF of NIC (n=12), RCVS (n=4) and PACNS (n=8) individuals is reported. PACNS CSF has measurable C5 in the CSF whereas RCVS and NIC were below detection. Horizontal bars represent the mean for each group. Dotted line represents lower limit of detection (LOD) for the assay (62 pg/mL) determined by standard curve with recombinant C5. Values reported represent values obtained through ELISA and multiplied by two to account for original sample dilution in 2X storage buffer (see Materials and Methods). Fisher’s exact test shows statistical significance at p=0.0036 for the comparison of PACNS to RCVS+NIC cohorts, using binary classifications of above or below LOD.

### Summary of findings

We have performed a comprehensive molecular analysis of CSF from patients with biopsy-proven PACNS. Among other potentially interesting proteins, we find enrichment in regulatory components of the alternative complement pathway, in addition to increased levels of the terminal component, complement C5. The activation state of C5, that is, whether or not C5 has been cleaved into C5 anaphylatoxin, remains to be determined. Irrespective, our results suggest that blocking activated C5 by small molecule inhibitors, including CCX168, warrants further investigation as a possible therapeutic intervention. Overall, our findings contribute to an expanded understanding the basis of neurological disease through identification of abnormal CSF protein fingerprints.

## Methods

### Study Approval

PACNS and NIC patients were recruited as part of a larger study analyzing biological samples from patients with suspected neuroinflammatory disease at UCSF. The UCSF Institutional Review Board (IRB) approved the study protocol, and participants or their surrogates provided written informed consent. RCVS controls were recruited as part of a larger study analyzing biological samples from patients with CNS vascular disorders at Cleveland Clinic. The Cleveland Clinic IRB approved the study protocol, and participants or their surrogates provided written informed consent.

### Experimental Procedures and Statistics

See Supplemental Materials and Methods.

## Author Contributions

Study design: CMB, MRW, JLD, JMG, RAH and LHC. Experiments: CMB, HR unless otherwise noted. Text: CMB. Statistical analysis: HR. Figures and tables: HR and MRW. Mass spectrometry data acquisition and analysis: GMK, AY. Patient care and clinical reports: HAS, KCZ, TT, RAH, LHC, MPG, JMC, AGS, JFM, SAJ, VCD, JMG and MRW.

## Acknowledgements

This study was supported by the UCSF Center for Next-Gen Precision Diagnostics supported by the Sandler Foundation and William K. Bowes, Jr. Foundation (C.M.B, H.R., J.L.D., M.R.W., J.M.G., H.A.S., K.C.Z.); UCSF Medical Scientist Training Program (H.R.); the Rachleff Foundation (M.R.W.), Chan Zuckerberg Biohub (J.L.D.); and NIH National Institutes for Neurological Disorders and Stroke (award number K08NS096117, M.R.W.). Its contents are solely the responsibility of the authors and do not necessarily represent the official views of the NIH. Special thanks to Dr. HajjAli and Dr. Calabrese (Cleveland Clinic, OH) for their scientific input throughout this process and for elevating the study through suggestion and sharing of RCVS controls. Mass spectrometry analysis was performed in the UCSF Mass Spectrometry Facility (A. L. Burlingame, director) supported by the Adelson Medical Research Foundation. We would like to thank Ms. Anna Karydas and Drs. Zachary Miller and Howard Rosen for contributing the NIC CSF samples to this study.

## Notes

**Conflict of interest statement** Dr. Gelfand reports no relevant conflicts of interest to the submitted work. Outside of the submitted work: Personal compensation for consulting from Biogen and Merck. Research support from Genentech, Quest Diagnostics, MedDay. Personal compensation for medical legal consulting. No other conflicts of interest exist.

